# PoxiPred: An artificial intelligence-based method for the prediction of potential antigens and epitopes to accelerate vaccine development efforts against poxviruses

**DOI:** 10.1101/2023.08.04.551871

**Authors:** Gustavo Sganzerla Martinez, Mansi Dutt, Anuj Kumar, David J. Kelvin

**Affiliations:** Department of Microbiology and Immunology, Faculty of Medicine, Dalhousie University, Halifax, Canada; Department of Pediatrics, IWK Health Center, Canadian Centre for Vaccinology CCfV, Halifax, Canada

**Keywords:** Poxviruses, Artificial Intelligence, Epitopes, T-cell

## Abstract

**Motivation:** *Poxviridae* is a family of large, complex, enveloped, and double-stranded DNA viruses. Members of this family are ubiquitous and well known to cause contagious diseases in humans and other types of animals as well. Despite significant progress in Artificial intelligence (AI) based methods, limited methods are available to predict the epitopes. In this study, we have proposed a unique method to predict the potential antigens and T-cell epitopes for multiple poxviruses.

**Results:** With PoxiPred, we developed an AI-based tool that was trained and tested with antigens and epitopes of *Poxviruses*. Our tool was able to locate 1,675 antigen proteins from 25 distinct *poxviruses*. From these antigenic proteins, PoxiPred also located 6,579 T-cell epitopes. PoxiPred is able to, on a single run, identify antigens and T-cell epitopes for poxviruses with one single input, i.e., the proteome file of any poxvirus.

**Availability:** The code that implements PoxiPred and the predicted antigens/T-cell epitopes is publicly available at https://github.com/gustavsganzerla/poxipred.

## 1 Introduction

Poxviruses (members of the *Poxviridae* family) are a family of double-stranded DNA viruses with genomes of approximately 135 to 360 kbp (Günther et al., 2017). Members of the large family exist throughout the world and can cause a plethora of diseases in humans and animals (Buller and Palumbo, 1991). Based on their genome architecture and evolutionary relationship, members of *Poxviridae* are classified into two subfamilies, namely, *Chordopoxvirinae* and *Entomopoxvirinae*. The subfamily *Chordopoxvirinae* is further divided into 18 genera; among them, four genera, *Molluscipoxvirus, Orthopoxvirus, Parapoxvirus*, and *Yatapoxvirus*, are known to cause human infections (Hughes et al., 2010; https://ictv.global/report/chapter/poxviridae/poxviridae). Popular contagious diseases associated with this subfamily include Mpox (formerly Monkeypox), smallpox, cowpox, and lumpy skin disease, which affects cattle (Dutt et al., 2023).

Epitopes, also known as antigenic determinants, are defined as the portion of foreign protein or antigen that could elicit an immune response mediated by antibodies, T, or B cells. Epitopes offer a targeted approach to vaccines against infectious diseases (Yang et al., 2021).

Recent advancements in Machine Learning (ML) methods have enabled the computational biologist-immunologist to design accurate epitopes. In these regards, viral proteins and peptides are commonly coded into numeric features such as Z-descriptors (Hellberg *et* al, 1897), representing their structural conformations in a way that ML applications can classify proteins/peptides with different functions (i.e., antigens and non-antigens; epitopes and non-epitopes, among others). Moreover, some of the methods that enable immunoinformatics have not yet fully adapted to the arrival of ML and still employ mechanistic statistics as a basis of classification (Doytchinova and Flower, 2007), which might fail to capture (Sganzerla Martinez et al, 2022) the distinctive signal portrayed by data with known immune function. We also argue that current methods are still limited to designing genome-specific epitopes to cope with emerging and re-emerging viral diseases. In the present study, we attempted to develop an AI-based method for predicting antigen proteins as well as designing T-cell epitopes targeting 25 poxviruses belonging to different genera of the *Chordopoxvirinae* subfamily.

## 2 Methods

We obtained 978 known T-cell epitopes from *orthopoxviruses*, including *Variola, Vaccinia, Cowpox, Mpox*, and *Ectromelia* viruses (Grifoni *et* al, 2022). The length of the epitopes ranges from 11±3 amino acids (aa). To obtain non-epitope data, we randomly selected a same-sized peptide in the same protein sequence. From the known epitopes, we mapped them back to their proteins of origin to obtain a dataset of antigen proteins, which resulted in 217 unique proteins spread across five *Orthopoxviruses*. To obtain non-antigen proteins, we randomly selected 217 non-antigen proteins in the proteome file of each virus.

We trained/tested two deep-learning Artificial Neural Networks (ANNs), the first for predicting antigen proteins and the second for predicting epitopes from the antigen proteins through a Python (version 3.9.7) script implementing the TensorFlow (version 2.12.0) package. Each ANN had its training/testing process cross-validated 10 times. Each protein sequence (antigen and epitope) was annotated by Z-descriptors (Hellberg *et* al 1987), converted to a same-sized 45-length vector by an Automated Cross Covariance (ACC) transformation (Yang *et* al, 2021), and normalized through the StandardScaler method found in the Scikit-learn package (version 1.3.0). We iterated over one, two, and three hidden layers, each with 10, 25, and 50 neurons to find the best set of hyperparameters. We stopped adding hidden layers and neurons to the models once they reached a test loss of < 0.1. For the antigen model, 500 learning epochs were used while the epitope model was given 100 epochs. Each model had its performance assessed by averaging the accuracy, precision, recall, specificity, and loss obtained at each fold of the train and test processes. In each ANN model, we identified the best threshold applied to the sigmoid function. Once the ANN was trained/tested satisfactorily per the accuracy, precision, recall, specificity, and loss of both train and test iterations, we obtained proteomes of the 25 poxviruses belonging to eight different genera of the subfamily *Chordopoxvirinae* including *Avipoxvirus* (03 members), *Capripoxvirus* (03 members), *Leporipoxvirus* (01 member), *Molluscipoxvirus* (01 member), *Orthopoxvirus* (09 members), *Parapoxvirus* (05 members), *Suipoxvirus* (01 member), and *Yatapoxvirus* (02 members), which were then downloaded from UniProt (https://www.uniprot.org/) in fast format. A detailed description of the species, proteome accessions, and number of proteins can be found in **Suppl. Table 1**.

Each protein in the proteome data was submitted to the antigen classifier. Only if a protein was predicted as having antigenic activity, up to five epitopes were attempted to be found. We considered the length of each predicted epitope ranging from 11±3. The epitope ANN would run until 5 epitopes were found or after all combinations of linear epitopes were exhausted. The resulting predictions consisted of *i*) the protein sequence with predicted antigenic activity; *ii*) all the epitopes (up to five) found; and *iii*) the function of the antigen protein. The Python code that implements the Poxviruses Immune Predictor (PoxiPred) is freely available at https://github.com/gustavsganzerla/poxipred.

## 3 Results

An overview of the entire pipeline constructed for the PoxiPred method is available in Fig. 1.

**Fig. 1.**
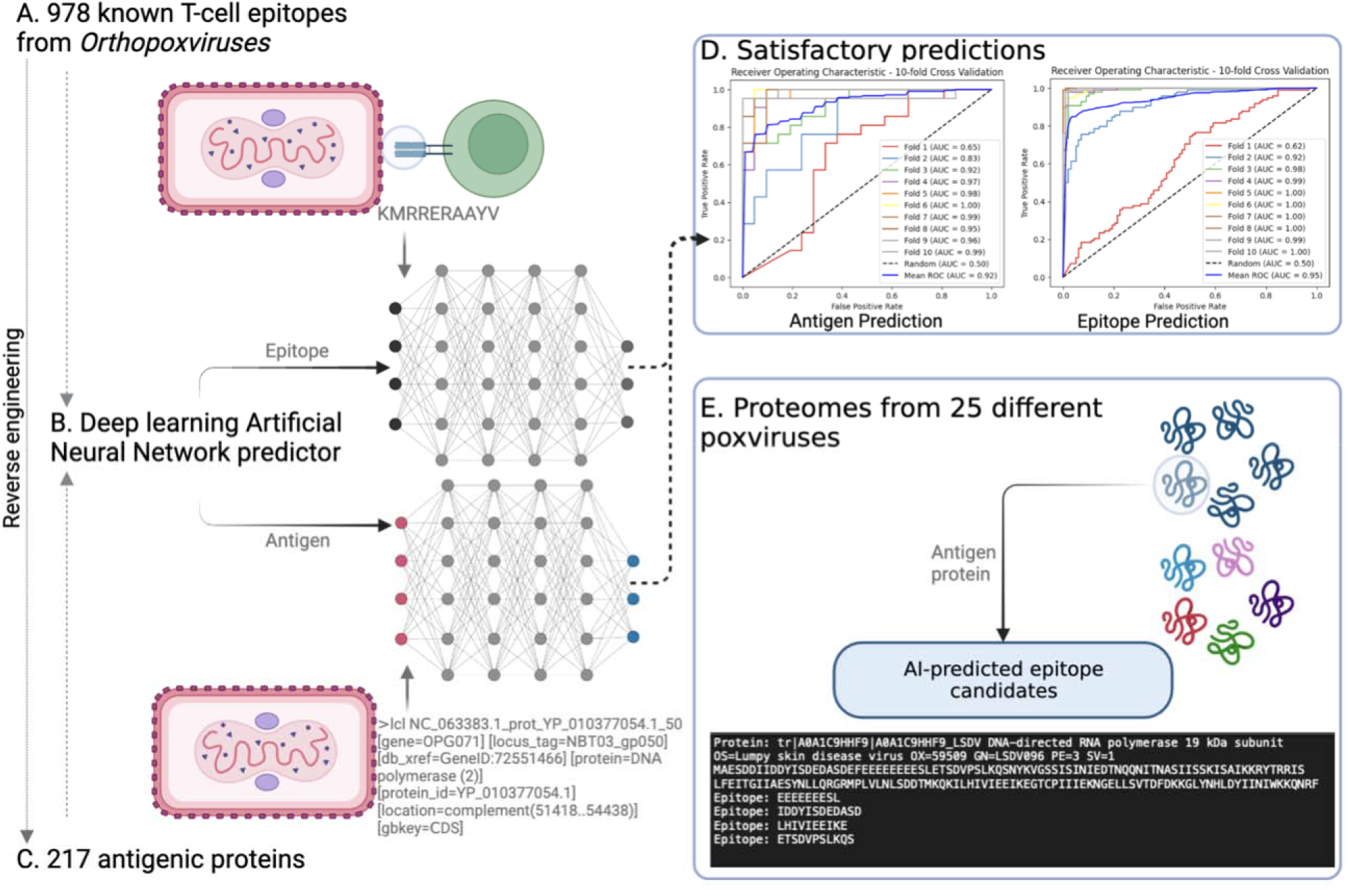
A flowchart of the PoxiPred method pipeline.

In Fig. 1-A, we depict the data with known epitopes across 5 different *orthopoxviruses*. Each of these epitopes was mapped back to their protein of origin (Fig. 1-C) to generate a dataset with known antigens. Both antigens and epitopes were classified with two separate deep-learning Artificial Neural Networks (Fig. 1-B), whose predictions resulted in an averaged Area Under the Curve (AUC) score of 0.92 and 0.95 (for antigen and epitope prediction, respectively) over a 10-fold cross-validation process (Fig. 1-D). Next, in Fig. 1-E, we selected the proteome files of 25 different poxviruses to predict their potential antigen proteins and their T-cell epitopes.

First, to predict the antigenicity of viral proteins, we ran an ANN trained/tested with 217 antigens from *orthopoxviruses*. We tuned the hyperparameters of the ANN to find that the best configuration would be three hidden layers with 50 neurons each trained over 500 epochs. This ANN model resulted in satisfactory test performance averaged over a 10-fold cross-validation process (accuracy = 95.9%, precision = 99.5%; recall = 92.3%; specificity = 99.5%, and loss = 0.06). We also identified the best limit applied to the output of the sigmoid function that would classify true positive and true negative antigens as 0.9999401. Moreover, to predict potential epitopes in the antigen protein, we ran an additional ANN trained/tested with 978 known epitopes from different *orthopoxviruses*. We tuned the hyperparameters of the ANN to find the best configuration as being three hidden layers with 50 neurons each trained over 100 epochs. This ANN resulted in satisfactory test performance averaged over a 10-fold cross-validation process (accuracy = 93.1%; precision = 99.6%; recall = 86.5%; specificity = 99.6%; and loss = 0.09). The results of this ANN were obtained by setting 0.9990533 as the threshold applied to the output of the sigmoid function in the output layer of the model. The result of each combination of hyperparameters for both the antigenicity and epitope ANN is available in **Suppl. Table 2**.

Upon the successful training/testing of the antigen and epitope prediction (mean AUC = 0.92 and 0.95, respectively) (Fig. 1-E), we fed our model with 4,517 proteins found in the proteome of 25 *poxviruses*. First, each protein was predicted as being an antigen or not. For these, we identified in total 1,675 proteins as antigens. For each protein predicted by PoxiPred as an antigen, we ran our T-cell epitope predictor to find up to 5 T-cell epitopes per antigen. In total, we obtained 6,579 *in-silico* predicted epitopes. The breakdown of the number of antigenic proteins and the epitopes predicted by the virus is available in **Suppl. Table 3**. Moreover, the CSV files containing the predictions are publicly available at https://github.com/gustavsganzerla/poxipred, following the columns: *i*) the aa sequence of the protein; *ii*) the predicted T-cell epitopes (up to 5 columns); and *iii*) the description of the protein predicted as an antigen.

Currently, the T-cell epitopes predicted by PoxiPred are derived from extensive ML approaches. There is a need to apply streamlined filters to the resulting epitopes such the epitope antigenicity, toxicity, allergenicity, and solubility. Also, other immunological factors such as the binding of the epitopes to T cell receptors and HLA diversity will be considered in future iterations. Moreover, PoxiPred might benefit from a user interface in which novel proteins of different poxviruses can map potential epitopes; currently, the team behind PoxiPred is addressing both questions as well as the inclusion of B-cell epitopes for future iterations. In the meantime, we believe that providing a set of in-silico predicted T-cell epitopes might assist scientists interested in immune responses against poxviruses, as in a single run users can predict antigen proteins and their potential T-cell epitopes as candidates for rapid vaccine development of emerging poxviruses.

## Acknowledgements

The corresponding author (DJK) is the Canada Research Chair in Translational Vaccinology and Inflammation.

