## Supplementary Table1 for "PoxiPred: An artificial intelligence-based method for the prediction of potential antigens and epitopes to accelerate vaccine development efforts against poxviruses"

| **S. No.** | **Name of the Virus** | **Proteome ID** | **Number of Proteins** |
| --- | --- | --- | --- |
| 1 | Lumpy skin Disease virus | UP000315615 | 156 |
| 2 | Myxoma virus | UP000291359 | 158 |
| 3 | Orf virus | UP000693994 | 130 |
| 4 | Sealpox virus | UP000202998 | 119 |
| 5 | Pseudocowpox virus | UP000117145 | 125 |
| 6 | Sheeppox virus | UP000318262 | 129 |
| 7 | Ectromelia virus | UP000130118 | 180 |
| 8 | Squirrel pox virus | UP000144311 | 141 |
| 9 | Camelpox virus | UP000107153 | 261 |
| 10 | Yaba monkey tumor virus | UP000008596 | 140 |
| 11 | Tanapox virus | UP000099606 | 155 |
| 12 | Swinepox virus | UP000000871 | 146 |
| 13 | Turkeypox virus | UP000142477 | 170 |
| 14 | Taterapox virus | UP000139570 | 220 |
| 15 | Vaccinia virus | UP000000344 | 218 |
| 16 | Variola Virus | UP000002060 | 198 |
| 17 | Cowpox virus | UP000097203 | 214 |
| 18 | Horsepox virus | UP000111173 | 228 |
| 19 | Monkeypox virus | UP000516359 | 183 |
| 20 | Molluscum contagiosum virus subtype | UP000000869 | 163 |
| 21 | Fowlpox virus | UP000150838 | 232 |
| 22 | Bovine papular stomatitis virus | UP000104372 | 130 |
| 23 | Canarypox virus | UP000168164 | 322 |
| 24 | Volepox virus | UP000203649 | 204 |
| 25 | Goatpox virus | UP000134642 | 149 |

**Table: - List of 25 poxviruses along with proteome IDs and number of proteins**
